# Rapid invariant encoding of scene layout in human OPA

**DOI:** 10.1101/577064

**Authors:** Linda Henriksson, Marieke Mur, Nikolaus Kriegeskorte

## Abstract

Successful visual navigation requires a sense of the geometry of the local environment. How do our brains extract this information from retinal images? Here we visually presented scenes with all possible combinations of five scene-bounding elements (left, right and back wall, ceiling, floor) to human subjects during functional magnetic resonance imaging (fMRI) and magnetoencephalography (MEG). The fMRI response patterns in the scene-responsive occipital place area (OPA) reflected scene layout with invariance to changes in surface texture. This result contrasted sharply with the primary visual cortex (V1), which reflected low-level image features of the stimuli, and parahippocampal place area (PPA), which showed better texture than layout decoding. MEG indicated that the texture-invariant scene-layout representation is computed from visual input within ~100 ms, suggesting a rapid computational mechanism. Taken together, these results suggest that the cortical representation underlying our instant sense of the environmental geometry is located in OPA.

## INTRODUCTION

Animals move around in their environments with grace and foresight, avoiding collisions with obstacles by charting viable paths based on their vision. This behaviour requires an animal’s visual system to provide its navigational circuits with information about the local environmental geometry. The human cortex contains visual areas that preferentially respond to visually presented scenes as compared to other stimuli, such as faces or objects. These areas include the parahippocampal place area (PPA; Epstein and Kanwisher, 1998) and the occipital place area (OPA; Grill-Spector, 2003; Dilks et al., 2013). PPA and OPA likely play a role in connecting visual perception with navigation, but their differential computational roles have not been fully established.

Boundaries of open spaces, such as walls, constrain navigation and are therefore an essential aspect of the environmental geometry that our brains must represent (for a review, see Brunec et al., 2018). Even small children automatically use room geometry to reorient themselves (for a review, see Spelke et al., 2010). Recent neuroimaging studies suggest a role for human OPA in detecting navigationally important cues from visual scenes. A functional magnetic resonance imaging (fMRI) study demonstrated that OPA encodes possible paths in a visual scene (Bonner and Epstein, 2017), and if processing in the OPA is temporarily disrupted using transcranial magnetic stimulation (TMS), a person’s ability to use boundaries in a navigational task is impaired (Julian et al., 2016).

Early fMRI studies already reported that the spatial layout, and not the presence of objects within the scene drives the scene-selective areas (Epstein and Kanwisher, 1998). Subsequent neuroimaging studies have made a distinction between open and closed sceneries (Harel et al., 2012; Kravitz et al., 2011; Park et al., 2011) and revealed the relevance of the vertical height of boundaries (Ferrara and Park, 2016). However, exactly how scene-selective areas represent the geometry of the local environment has not been established.

Here we ask how individual scene-bounding elements and their compositions are represented in scene-selective cortical areas. We test different brain regions for an explicit representation of the 3D geometry that is invariant to surface appearance. We created a novel set of synthetic scene stimuli, in which we systematically vary the spatial layout of the scene by switching on and off each of five spatial boundaries (three walls, floor, and ceiling; see Fig. 1). The resulting 2^5^ = 32 layouts are rendered in three different styles of surface appearance (empty room, fences, urban space), yielding 96 scene stimuli. These stimuli were presented to 22 subjects, each of whom participated in both an fMRI and a magnetoencephalography (MEG) experiment. Whereas fMRI provides sufficiently high spatial resolution to resolve representations within a given brain region, MEG provides millisecond temporal resolution, enabling us to track the dynamics of cortical processing (Carlson et al., 2013; for a review, see Hari and Salmelin, 2012). We investigated to what extent OPA and PPA encode the scene layout in terms of the presence and absence of the scene-bounding elements, and how rapidly the respective representations emerge following stimulus onset.

**Figure 1.**
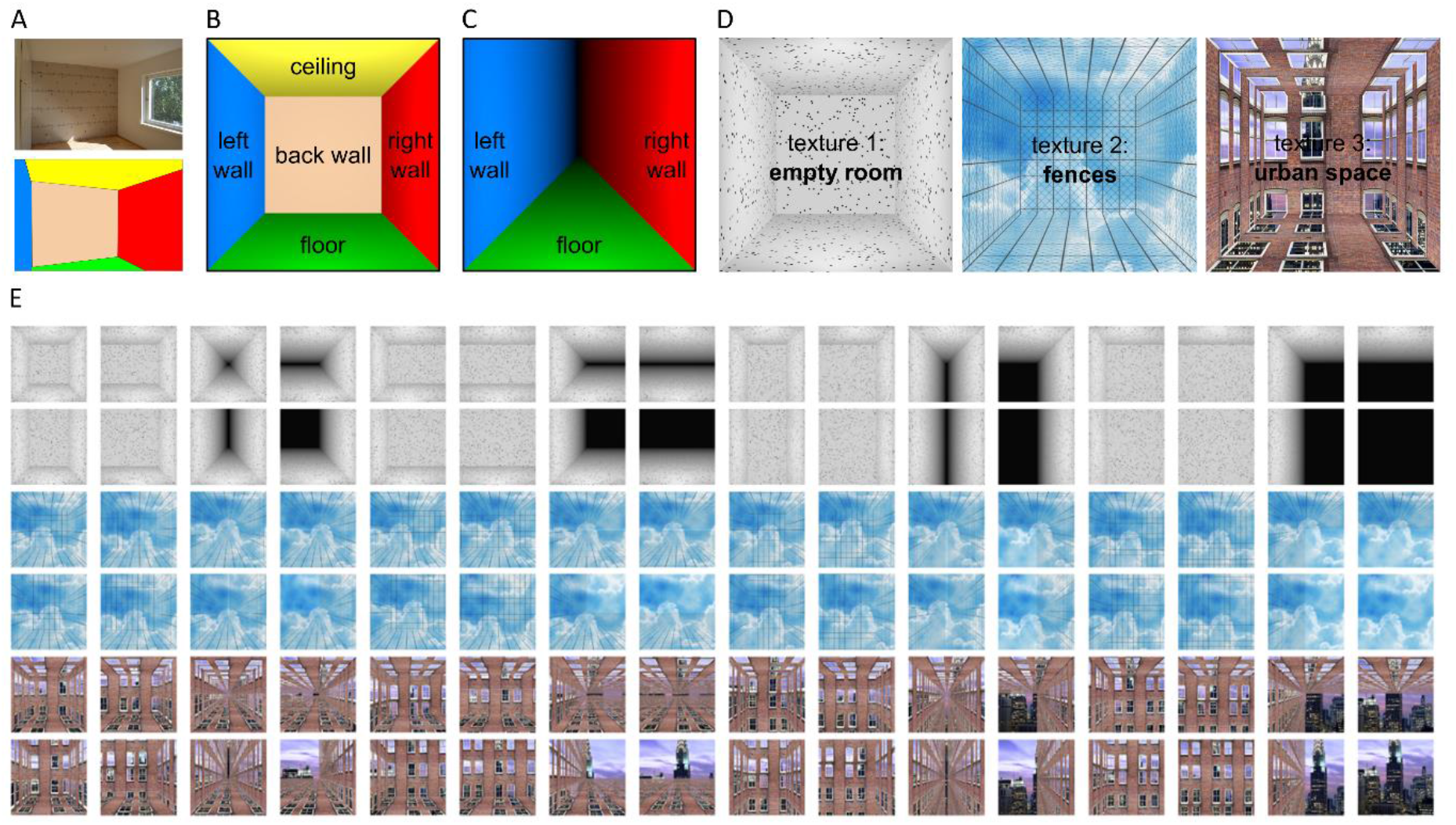
Stimuli to test the hypothesis that human scene-responsive cortical areas encode scene layout. A) The spatial layout of a room is captured by the fixed scene-bounding elements, such as the walls. B) We created a complete set of spatial layouts using 3D modelling software by switching on and off the five bounding elements: left wall, back wall, right wall, floor, and ceiling. C) For example, by switching off the back wall and the ceiling, we create a long, canyon-like environment. D) Textures and background images were added to the scenes to enable us to discern layout representations from low-level visual representations. E) The complete set of scenes included 32 different spatial layouts in 3 different textures, resulting in 96 scene stimuli.

## RESULTS

### OPA discriminates layouts better than textures whereas the opposite is true for PPA

We measured fMRI responses to the 96 different scene images (32 layouts in each of three different textures, Fig. 1E), while subjects fixated the stimuli centrally. Subjects were instructed to pay attention to the layout of the scene elements. Occasionally, the stimulus image was followed by an arrow pointing to one of five possible directions and the subject’s task was to tell with a button press whether the preceding layout had a bounding scene-element in that direction (*e.g*., an arrow pointing to the left would prompt the subject to report whether the left wall was present in the previous scene). Four regions of interest (ROIs) were defined based on criteria independent of the main experiment: primary visual cortex (V1) was defined on the basis of cortical sulci (Hinds et al., 2008), and OPA, PPA and retrosplenial cortex (RSC) were defined on the basis of functional localizer data using a different set of scenes, faces, objects, and textures as stimuli. Figure S1 shows the average responses separately for each stimulus in each ROI. In V1, the texture affected the response strength (empty < fence < urban), likely reflecting the amount of low-level image detail in the stimulus images. In OPA and PPA, the difference in response strength between the stimuli was smaller than in V1. In RSC, many of the stimuli did not evoke a measurable response, and hence, results for RSC are only shown in Supplementary Figures.

First, we asked whether we can discriminate the fMRI response patterns evoked by the different scene stimuli. The discriminability of each pair of stimuli was evaluated by fitting a Fisher linear discriminant (Nili et al., 2014) to the response patterns from half of the fMRI data and by testing the performance on the response patterns from the other half of the fMRI data (split-half cross-validation). The analyses were done on individual data and the results were pooled across the 22 subjects. First, we evaluated whether we can better discriminate the layout or the texture of a scene from the response patterns in different ROIs. Figure 2 shows the average linear-discriminant *t* (LD*t*) values for scenes that differ in layout (gray bars) and for scenes that differ in surface texture (black bars). In V1 and PPA, the LD*t* values were higher for texture than layout discrimination. Moreover, Figure S2 shows that in PPA, but not in V1, the scene discriminability was consistently higher when both the layout and the texture were different compared to scenes that only differed in their layout. As the texture defines the scene’s identity, these results suggest that PPA is involved in texture-based scene categorization rather than representing scene layout. In contrast to the results in PPA, in OPA the average discriminability was higher between scenes that differ in layout than in texture (Figure 2).

**Figure 2.**
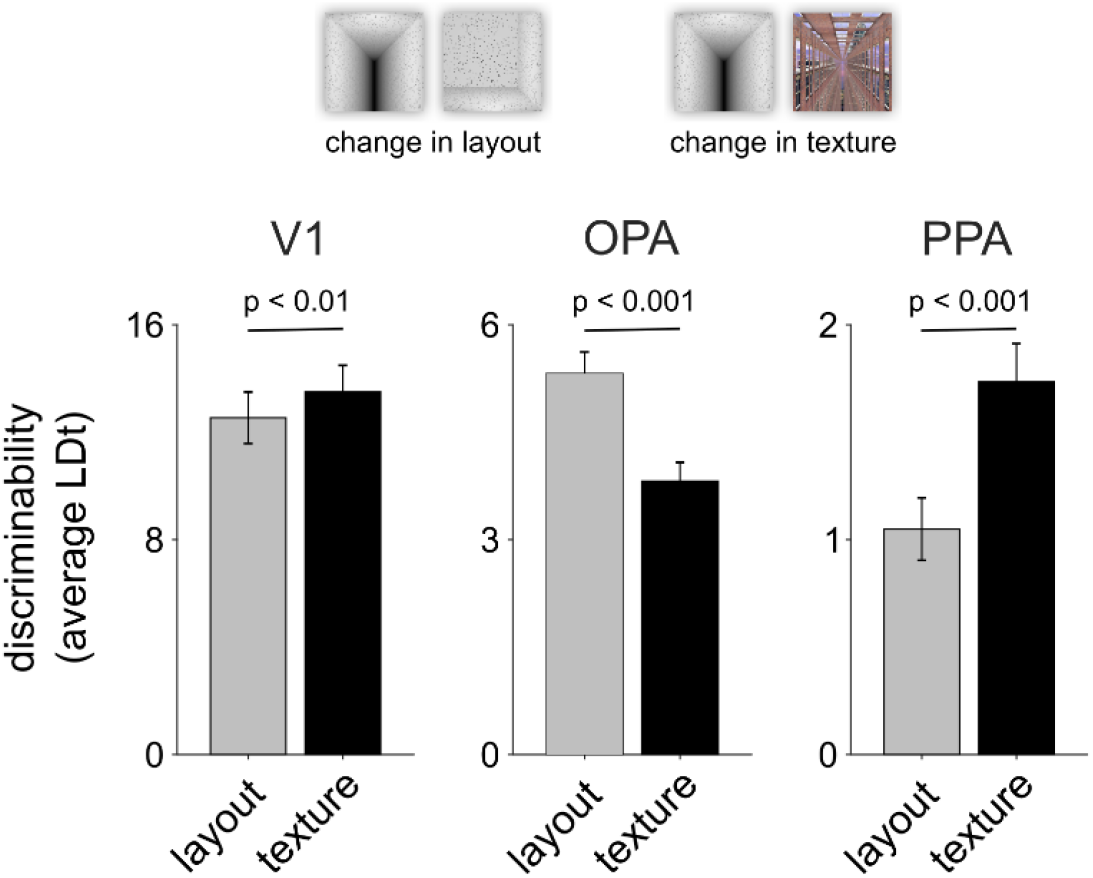
Layout vs. texture decoding. The average discriminability across all scene pairs that differ in the layout but are of the same texture (layout decoding; grey bars) and all scene pairs that have the same layout but differ in the texture (texture decoding; black bars) are shown separately for V1, OPA and PPA. In V1 and PPA, a change in the texture had on average a larger effect on the discriminability of the fMRI response-patterns than a change in the layout, whereas the opposite was true for OPA. The p-values are from two-tailed signed-rank tests across the 22 subjects; the error bars indicate standard errors of the mean (SEMs) across the subjects. See Figure S1 for average fMRI responses for all 96 individual stimuli for each region-of-interest, and Figure S2 for fMRI response-pattern discriminability separately for each stimulus pair.

### Layout discrimination in OPA generalizes across surface-textures

In order to test whether any of the regions contain a layout representation that is invariant to texture, we fit the linear discriminant to the response patterns for a pair of layouts in one texture and tested its performance with the same layouts in another texture. Successful cross-decoding across textures would suggest a scene-layout representation that is tolerant to a change in the surface-texture, and at the same time, rule out confounding effects of low-level image-feature differences on layout discrimination. Figure 3A-B shows a schematic illustration of the analysis. The distinctiveness of each layout pair is shown separately (diagonal matrices) as well as the ability of the discriminants to generalize to other surface-textures (off-diagonal matrices). These matrices will be referred to as representational dissimilarity matrices (RDMs).

**Figure 3.**
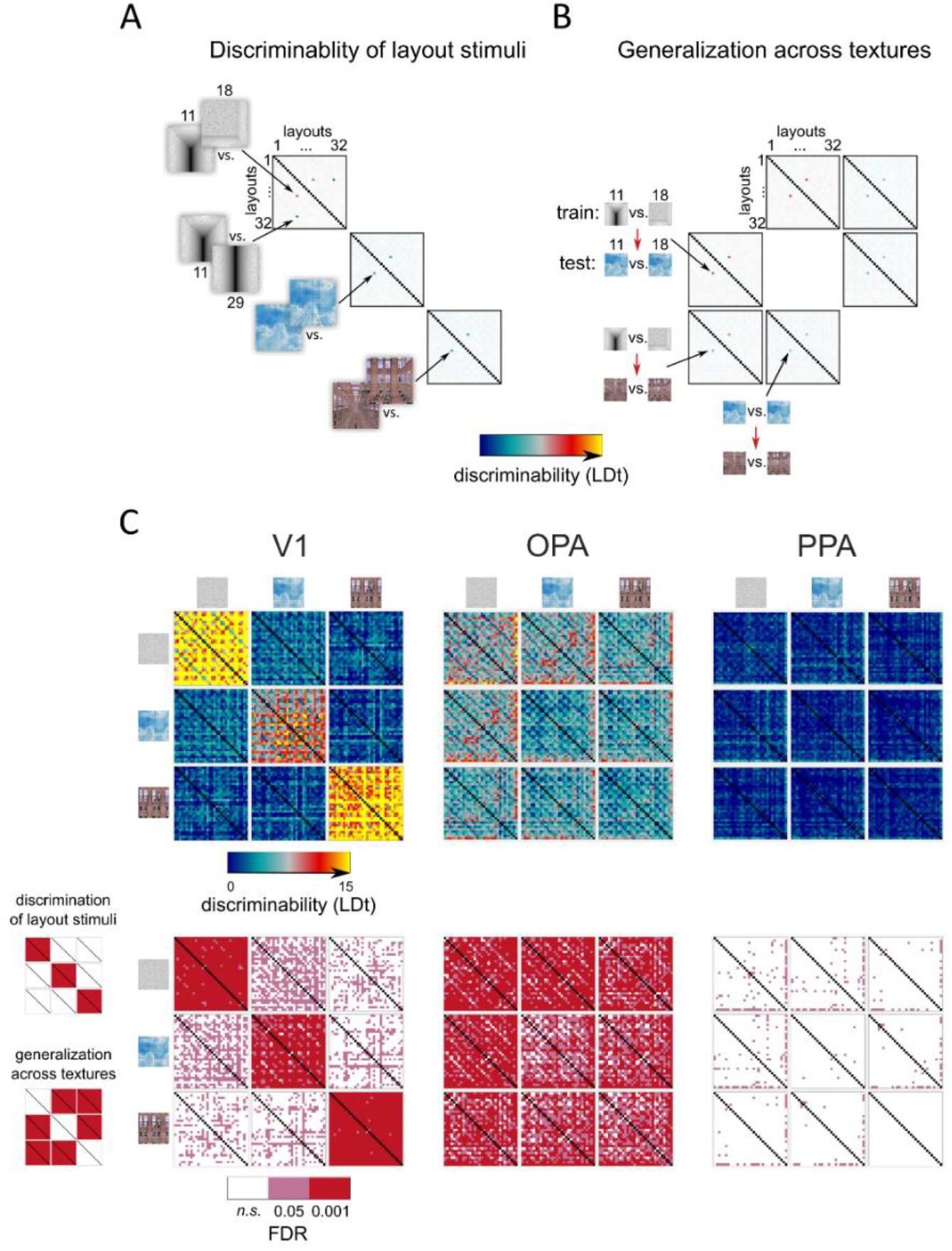
Discriminability of the fMRI response-patterns for each pair of layout stimuli in V1, OPA and PPA and the generalization of the result across textures. A) The discriminability of the layout stimuli from the fMRI response-patterns was evaluated for each pair of the 32 layouts, separately for the three textures. The Fisher linear discriminant was fitted to response-patterns obtained from training data (odd fMRI runs) and the performance was evaluated on independent testing data (even fMRI runs). The results are shown as linear discriminant t-values (LDt; Nili et al., 2014; Walther et al., 2016). The analyses were done on individual data and the results were pooled across subjects. B) The generalization of the layout discrimination across different surface textures was evaluated by fitting the Fisher linear discriminant to response-patterns corresponding to a pair of layouts in one texture and evaluating the performance on the response-patterns corresponding to the same layout pair in another texture. All combinations of texture pairs were evaluated. A high LDt-value suggests successful generalization of layout discrimination across surface textures, and hence a layout representation that is tolerant to a change in surface texture. C) The upper row shows the average LDt values for all pairs of stimuli. Along the diagonal, the three matrices reflect the distinctiveness of the response-patterns between each pair of layout stimuli in the same texture. The six off-diagonal matrices show generalization across textures (training between two layouts in one texture, testing on the same pair in another texture). The bottom row shows the expected false-discovery rates (FDR; n.s., not significant); the p-values were obtained using a two-sided signed-rank test across 22 subjects. In V1, the spatial layout stimuli elicited distinct response-patterns but the result did not generalize across textures. In OPA, most layouts evoked a distinct response-pattern and the results generalized across textures. In PPA, only few of the layouts evoked significantly different response patterns.

Figure 3C shows that in V1 the layouts evoked distinct response patterns, but the results did not generalize across textures. This result suggests that the pattern-discriminability in V1 was due to confounding low-level image feature differences between the same-texture spatial layouts instead of an explicit representation of layout. In OPA, on the contrary, the discriminants for stimulus pairs generalized across textures, suggesting that OPA encodes the layout of a visual scene invariantly to manipulations of surface texture. Finally, although PPA responded to the stimuli (Figure S1) and its average layout discriminability was above chance (Figure 2), its patterns did not enable reliable discrimination of most pairs of layouts (Figure 3C).

Figure 4 shows the average LD*t*-RDMs and the corresponding false discovery rate matrices for V1, OPA and PPA, summarizing results from Figure 3. Corresponding multidimensional scaling (MDS) visualizations of the representational relationships are shown in Figure S3. The average RDMs and the MDS visualizations reveal that the presence of the back wall had a strong effect on the distinctiveness of the response patterns in V1, OPA, and PPA. In contrast to the other scene elements, the back wall covered a larger part of the visual field and was centred on the point of fixation. Given its larger retinal extent and the cortical magnification of the central region (Duncan and Boynton, 2003), the back wall had a much larger cortical representation than the other scene elements in early visual areas, especially in V1. In the scene-responsive regions, the added depth to the scenes by the removal of the back wall could also contribute to response-pattern discriminability. In OPA, we observe groupings that are consistent both in the within- and across-texture analysis (Figs. 4, S3). For example, pairs of scenes that only differed in the presence of the ceiling elicited similar response patterns (blue off-centre diagonal in the LD*t*-matrices), suggesting that the ceiling did not strongly contribute to the layout representation in OPA. Moreover, the number of the bounding elements present in the layout appears to have an effect on pattern distinctiveness (clusters in the MDS plots; Fig. S3).

**Figure 4.**
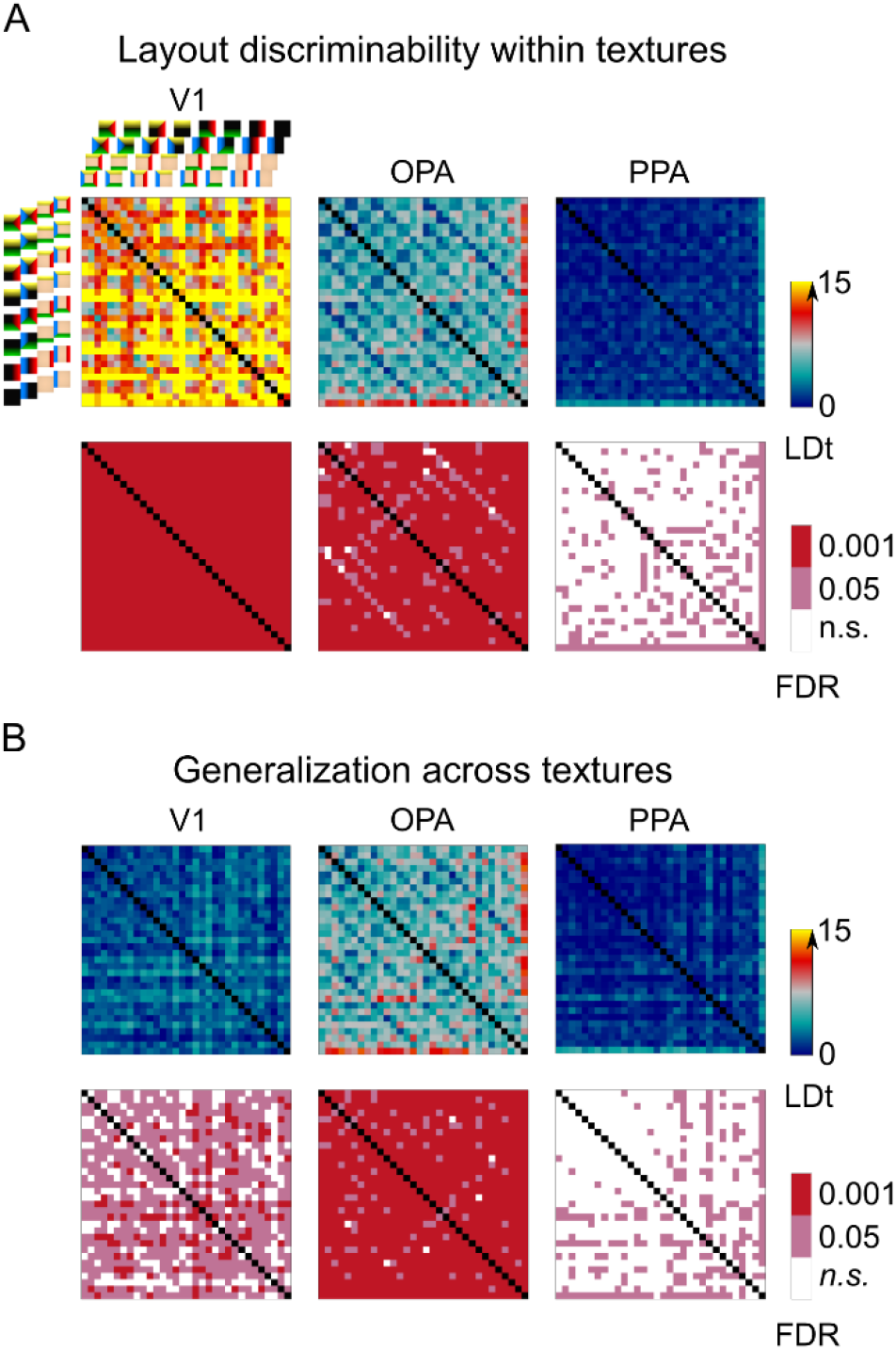
Representational geometry of scene layouts generalizes across textures in OPA. A) The distinctiveness of the fMRI response-patterns for the spatial layouts are shown as captured by the LDt values. The analyses done separately for the three textures were averaged (shown separately in Fig. 3). The lower row shows the corresponding false discovery rates (two-tailed signed-rank test across 22 subjects). B) The generalization performance of the discriminant across different textures was evaluated by fitting the linear discriminant to a pair of spatial layouts in one texture and testing the discriminant on the same pair of layouts in another texture. The analysis was done for each combination of textures and the results were averaged (shown separately in Fig. 3). The lower row shows the corresponding false discovery rates. In OPA, the representation shows texture-invariance. The multidimensional-scaling visualizations of the distinctiveness of the response-patterns are shown in Figure S3.

Although some layout-pairs could still be discriminated by the V1 response-patterns when the analysis was done across different textures (Fig. 4B), the overall decoding performance was significantly worse than when the analysis was done within the textures (Fig. 4A). In V1, but not in OPA, the representational geometry, as visualized using the MDS (Fig. S3), was also clearly different between the within-texture and cross-texture analyses. In PPA, the average discriminability of the layouts was low. OPA stands out against V1 and PPA in that it discriminated spatial layouts of the same texture and the result generalized across textures (middle panels in Fig. 4A and 4B). This finding is consistent with a texture-invariant scene-layout representation in OPA.

### The OPA representation is reflected in early MEG responses

All subjects participated also in an MEG experiment to characterize the temporal dynamics of scene-layout encoding. Similar to the fMRI experiments, the subjects fixated the stimuli centrally and were instructed to pay attention to the layouts. Occasionally, the stimulus image was followed by an arrow pointing to one of the five possible directions and the subject’s task was to tell with a finger lift whether the preceding layout had a scene-bounding element in that direction. Representational dissimilarity matrices were constructed based on the cross-validated Mahalanobis distance between the evoked responses of each pair of spatial layouts. For fMRI, the cross-validated distance estimator LD*t* was chosen based on previous results (Walther et al., 2016). For MEG, cross-validated distance estimators are also recommended (Guggenmos et al., 2018), but the effect of noise normalization is less well understood. We therefore used the crossvalidated Mahalanobis distance (also termed linear discriminant contrast, LDC; Walther et al., 2016), which omits the final noise normalization. Figure S4 shows a comparison of LD*t* and LDC RDM reliability in our data. The MEG-RDMs were compared to the fMRI-RDMs using Kendall’s tau-a rank correlation (Nili et al., 2014). The analyses were done on individual data, separately for each texture. Representational dissimilarities were then averaged across textures for each subject, and the significance testing treated subject as a random effect.

Figure 5A shows that, as expected, both V1 and OPA fMRI-RDMs correlated with the MEG-RDMs. To find out the unique contribution of the OPA to the correlation, MEG-RDMs were fitted as a linear combination of the V1 and OPA fMRI-RDMs. OPA showed a unique contribution to the MEG-RDM fit early after stimulus onset. The unique OPA contribution becomes significant at 60 ms after stimulus onset (two-sided signed-rank test across subjects, multiple testing accounted for by controlling the false discovery rate at .01) and peaks at about 100 ms (Fig. 5B). Importantly, only the OPA showed a significant match between fMRI and MEG across-texture generalization of layout discriminants (Fig. 5C). In other words, MEG reflected the surface-texture invariant representation of the spatial layouts similarly to the OPA fMRI-RDM, and the similarity emerged early in the MEG data (significant at 65 ms, peaking at about 100 ms). These results suggest an early, texture-invariant encoding of scene-layout in the OPA.

**Figure 5.**
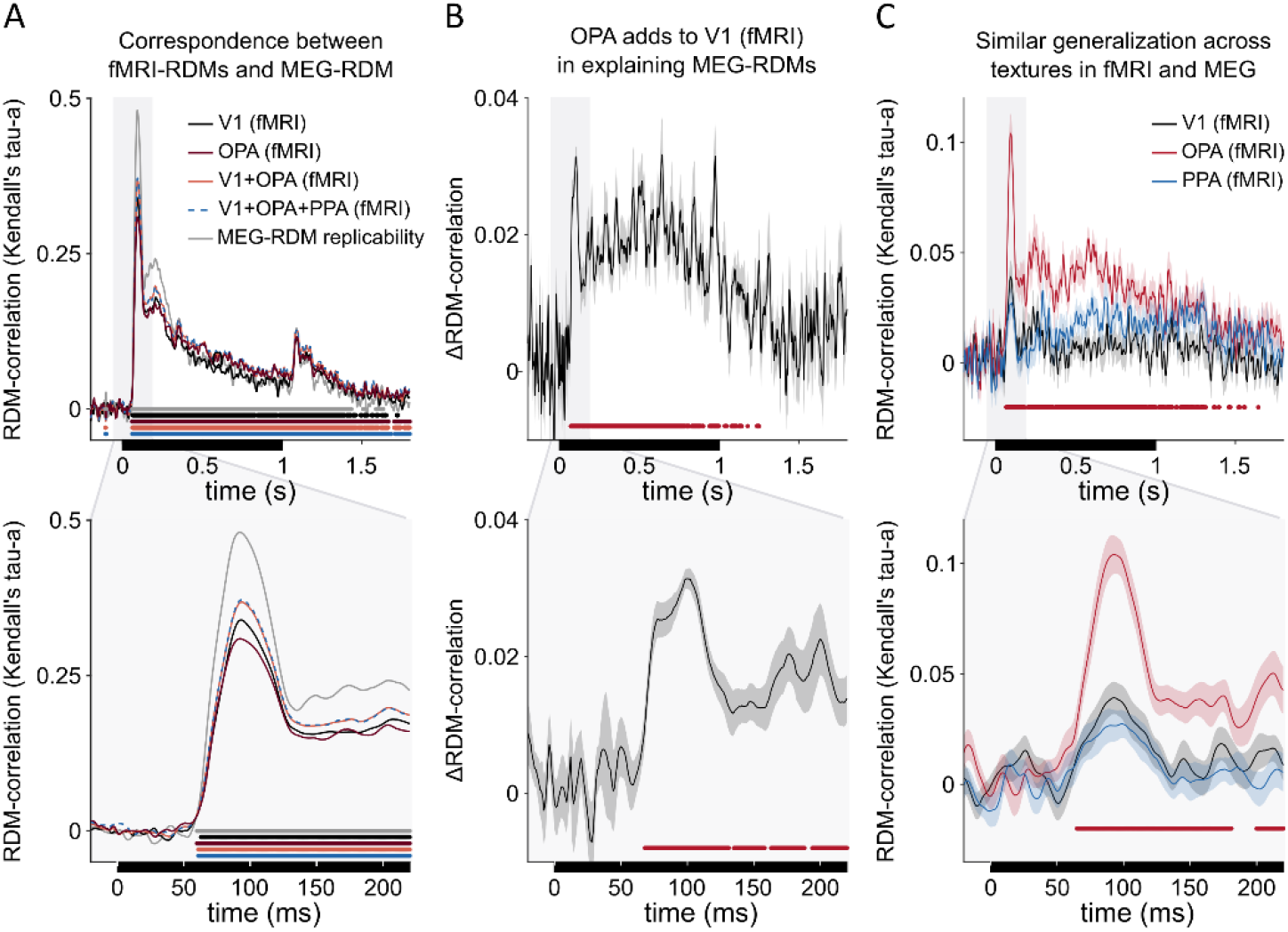
Early correspondence between the representations in OPA (fMRI) and MEG. A) The Kendall tau-a rank correlation between the MEG-RDMs and fMRI-RDMs are shown. The upper panel shows the full time-course (stimulus-on period indicated by black bar) and the lower panel highlights the early time-window. The black line shows the correlation between the MEG and V1-fMRI, and the dark red line shows the correspondence between the MEG and OPA-fMRI. The correspondence between the MEG-RDMs and cross-validated, fitted linear combinations of multiple fMRI-RDMs are also shown (light red line for V1 and OPA, blue dashed line for V1, OPA and PPA). The analyses were done separately for each subject and for each texture, and the results were averaged. Significant time points indicated with the thick lines (false-discovery rate, FDR, of 0.01; p-values computed with two-tailed signed-rank test across the 22 subjects; FDR adjusted across time-points). The grey line indicates the amount of replicable structure in the MEG-RDMs across subjects. Figure S4 evaluates the effect of the distance estimator for the reliability of the RDMs. B) The OPA significantly adds to V1 in explaining the MEG-RDMs already in the early time-window after stimulus onset. Shaded regions indicate SEM across subjects, red line indicates false-discovery rate of 0.01 (two-tailed signed-rank test across the 22 subjects; FDR across time points). C) Both MEG and fMRI -RDMs were also constructed based on how well the layout discrimination results generalized across surface-textures (see also Figs. 3–4). Only OPA-fMRI, not V1 or PPA, shows a significant correlation with MEG when the texture-generalized RDMs are compared (red line indicates false discovery rate of 0.01 between OPA-fMRI-RDM and MEG-RDM; two-tailed signed-rank test across the 22 subjects).

### Scene-bounding elements explain OPA representational geometry better than GIST

To characterize the geometries of the scene-layout representations, a set of models (Fig. 6) was fitted to the fMRI- and MEG-RDMs. We aimed to model both the contribution of the low-level image features and the presence of the scene-bounding elements. The GIST (Oliva and Torralba, 2001) model was included to capture low-level image feature differences between the scene stimuli. Separate GIST RDMs were constructed for each texture. The first scene-layout-based RDM model consisted of the Hamming distance between the binary vectors, indicating which scene-bounding elements were present and absent. For each pair of scenes, the Hamming distance is the number of discrepant scene-bounding elements. The Hamming distance assigns equal importance to all elements: vertical walls, floor, and ceiling. Hence, we name this model “ewalls” to stand for equally weighted walls (see top row in Fig. 6). In order to model the possibility that the brain representation does not weight all scene-bounding elements equally, we included a separate RDM component for each of the five scene-bounding elements and modelled the RDM as a fitted linear combination of the components (model “fwalls”, for fitted walls; middle row in Fig. 6). In addition, the number of walls present in a scene was included (model “nwalls”, for number of walls; top row in Fig. 6), predicting similar response patterns for two layouts with a similar number of walls. This model can be interpreted as reflecting the size of the space depicted in the scene (ranging from open to closed). For possible interaction effects between specific scene-elements (e.g., the presence of both the floor and the left wall forming an edge), interaction models were also constructed (two bottom rows in Fig. 6). The models were fitted using non-negative least squares (Khaligh-Razavi and Kriegeskorte, 2014) and tested by cross-validation across subjects.

**Figure 6.**
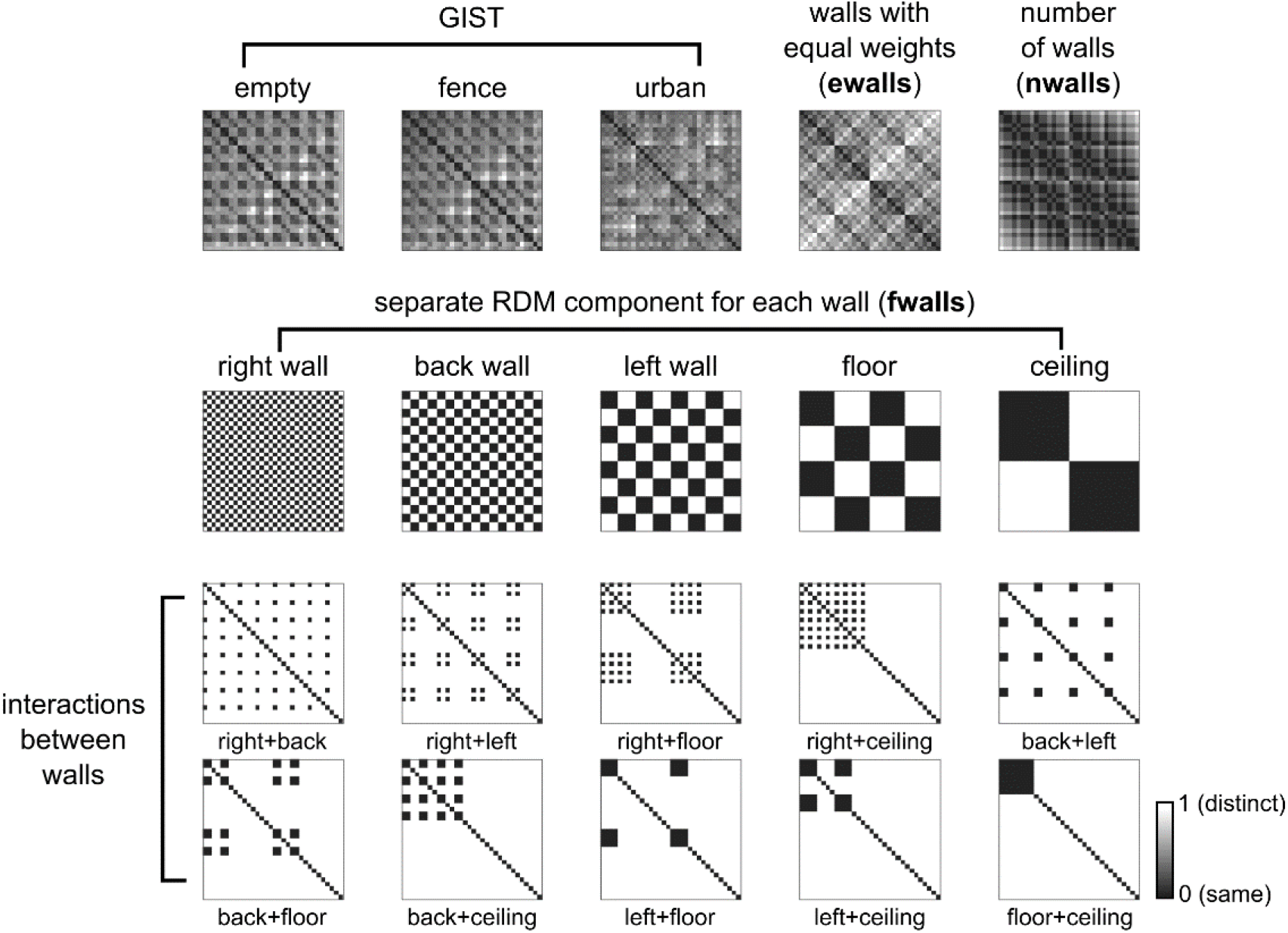
The representational geometries were modelled as linear combinations of representational components. The first three model-RDMs capture the GIST features (Oliva and Torralba, 2001) of the stimuli (low-level image feature similarity), shown separately for the three different textures. The next model predicts the responses for a scene-layout-based representation with equal contribution of all walls, calculated as the hamming distance between the layouts (percentage of same boundaries present in a pair of scenes; predicting a similar response to two scenes with same boundaries and distinct response for a pair of scenes with different boundaries). The last model on the top row is constructed based on the number of scene-bounding elements present in each scene (similar response to a pair of scenes with the same number of scene-elements present in the scene), roughly reflecting the size of the space. The middle row shows the five model-RDMs capturing separately the presence of each of the five scene-bounding elements in the spatial layouts. The last ten model-RDMs reflect the interactions between the walls.

The GIST model explained the V1 RDM (fMRI) better than any of the scene-layout models (nwalls, ewalls, fwalls; Fig. 7A–B). For OPA, results were strikingly different. The GIST model was still better than the number of walls (nwalls, reflecting the degree of openness) at explaining the OPA RDM (Fig. 7A–B) but there was no significant difference between the GIST model and the scene-layout model with equal contribution of each scene-bounding element (ewalls, Fig. 7A–B). Moreover, the fitted combination of scene-bounding elements model (fwalls) captured the representation significantly better than the GIST model (Fig. 7A–B). Adding the GIST component and the wall-specific RDM components in a fitted model did not increase explained variance in OPA (it did in V1). Explained variance in OPA was increased by adding the number-of-walls component, suggesting that scene openness is reflected the representation. Finally, adding components for the interactions between the scene-bounding elements increased the explained variance slightly in both V1 and OPA. In PPA, the best predictions were obtained by combining all the components (GIST, walls with fitted weights, number of walls). In contrast to V1 and OPA, however, adding the interaction components did not significantly improve predictions for PPA.

**Figure 7.**
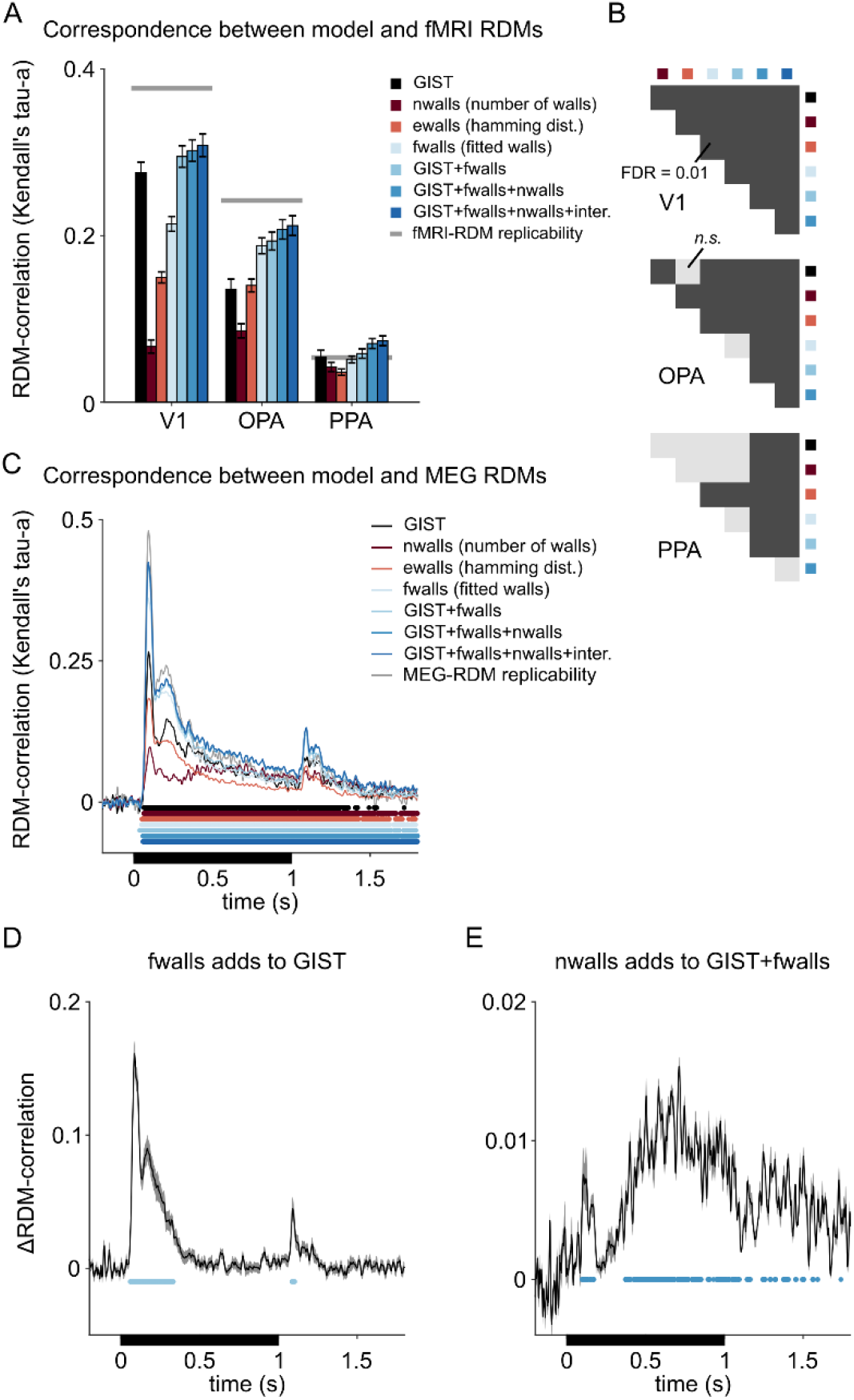
The contributions of low-level image differences and true scene-layout based representations in the fMRI- and MEG-RDMs. A) The mean Kendall’s tau-a correlation between the models and fMRI-RDMs are shown separately for V1, OPA and PPA. In V1, there is only a small improvement in the correlations when the GIST model is complemented with other models. In OPA, the fitted combination of scene-bounding elements model (fwalls) alone already explain the representation better than the GIST model. The grey lines indicate the amount of replicable structure in the fMRI-RDMs across subjects. The error bars indicate standard errors of the mean (SEMs) across the subjects. B) Pairwise comparisons between the models, shown separately for V1, OPA and PPA. The color-codes and order of the models is the same as in (A). Dark grey colour indicates significant difference between the models (two-tailed signed-rank tests across the 22 subjects, multiple testing accounted for by controlling the false discovery rate at .01), light grey indicates non-significance. C) The time-courses of the model-RDM fits to the MEG-RDMs are shown. The colours are the same as in (A). The grey line indicates the amount of replicable structure in the MEG-RDMs across subjects. The thick lines indicate significant correlations (false discovery rate of 0.01 based on two-tailed signed-rank tests across the 22 subjects). The black rectangle shows the timing of the stimulus. D) The mean increase in the Kendall’s tau a rank correlation is shown when the fitted wall-RDMs were included in the fitting compared to having only the GIST model. Time-points with significant increase are indicated with the blue lines (false discovery rate 0.01; two-tailed signed-rank test across subjects). E) Including the RDM based on the number of walls present in the scenes further increased the model fit, especially at the later timepoints (blue lines indicate false discovery rate 0.01; two-tailed signed-rank test across subjects).

Figure 7C shows the model crossvalidation results for the MEG RDMs, illustrating the dynamics of how the different models (Fig. 6) captured the representations. As expected, the GIST model captured the early representation (black line). However, including the scene-bounding elements improved the fit early. Figure 7D shows the unique contribution that scene-bounding elements added to the GIST model, peaking at about 100 ms after stimulus onset. Similar to the fMRI modelling, the contribution of each of the scene-bounding elements was modelled separately and the result was crossvalidated across subjects. Moreover, consistent with the fMRI results for OPA and PPA, the number of walls present in the scene also significantly contributed to the MEG-RDMs, especially at the later timepoints (Fig. 7E).

To further evaluate the contribution of each representational component, Figure 8 shows how much is gained in total explained variance by each of the model components. The analysis was done by leaving out each model component in turn and evaluating the proportion of variance explained between the full model and the reduced model. The interactions (two bottom rows in Fig. 6) were excluded from the full model as they did not significantly increase the adjusted explained variance in fMRI or MEG data (see Figs. S5–S6). The GIST model dominates the explained variance in V1. In both V1 and OPA (fMRI), the contribution of the back wall was stronger compared to the other walls, likely reflecting its larger visual field extent, and in OPA possibly also reflecting the added depth to the scenes by the removal of the back wall. In OPA, the floor had a significantly larger contribution than the right wall, the left wall, and the ceiling (two-sided signed-rank test across subjects, multiple testing accounted for by controlling the false discovery rate at .05). In the MEG results, corresponding results were found with the GIST model dominating the gain in the explained variance, but also a pronounced representation of the floor compared to right and left wall and the ceiling (bottom row in Fig. 8C), consistent with the results found in OPA (fMRI).

**Figure 8.**
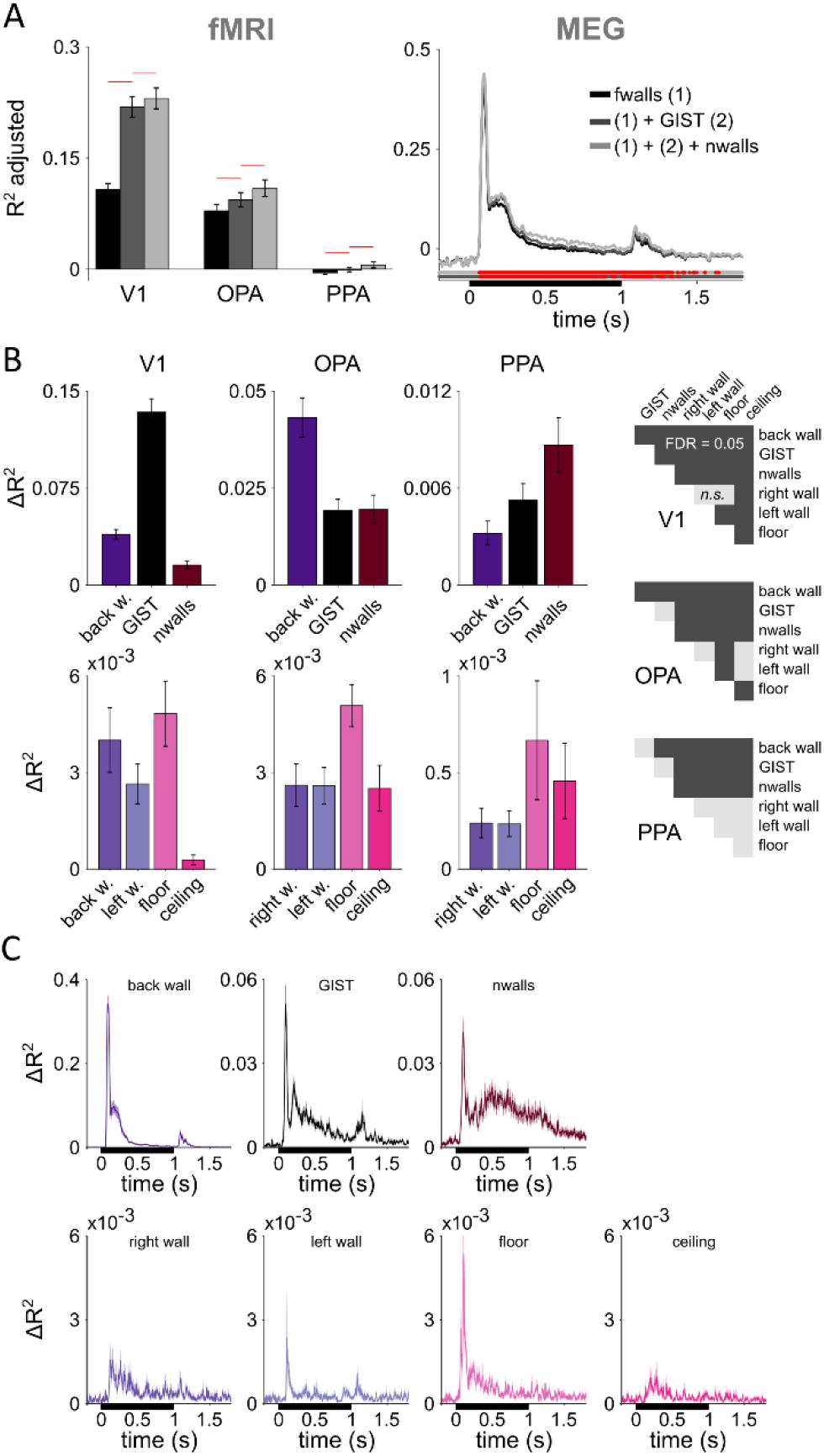
The unique variance explained by each model component. A) The explained variance (R^2^ adjusted) of the fitted joint models containing components for the five scene-bounding elements (black bars/line), components for the five scene-bounding elements and the GIST (dark grey bars/line), and components for the five scene-bounding elements, the GIST, and the number of walls (light grey bars/line). The red lines indicate a significant improvement compared to a model with fewer components (one-tailed signed-rank tests across the 22 subjects, multiple testing within a region-of-interest or across time-points accounted for by controlling the false discovery rate at .05). See Figures S5–6 for the results for each texture separately and including also the component models for interactions between walls (overall no further improvement in the adjusted explained variance). B) The gain in the variance explained by each of the model components was calculated by comparing the explained variance between the full model (light grey in A) with a model where this one component was left out from the joint fit. Each model component was left out in turn, and the results are shown separately for B) V1, OPA and PPA (fMRI) and C) MEG data. Error bars (fMRI) and shaded regions (MEG) are standard errors of the mean across subjects. In different regions-of-interest (fMRI), the different model components made different contributions to the representation; see the half-matrices for results from significance testing (two-tailed signed-rank tests across the 22 subjects, multiple testing within a region-of-interest accounted for by controlling the false discovery rate at .05). The pronounced contribution of the floor to the scene representations (fMRI-OPA and MEG) is consistent with the behavioural results shown in Figure S7.

### Fixations during free viewing favour the floor over the ceiling

Before the neuroimaging experiments, all 22 subjects participated in a behavioural experiment to familiarize themselves with the stimuli. The subjects were shown the 96 stimulus images (Fig. 1E) once in a random order. We tracked their eye gaze when they freely explored the scenes. The presentation time was short (1 second) and the subjects were asked to always bring their gaze back to the central fixation cross between the stimuli. Without any specific instructions on where or what to look at in the images, subjects made more saccades to the lower than upper visual field, and more specifically, to the floor than to the ceiling (Fig. S7; FDR = 0.0035; 0.0077; 0.003 for each texture separately; two-tailed signed-rank test across 22 subjects, FDR across the combinations of wall pairs). This result suggests that attention is automatically directed to the lower parts of a scene, where the navigationally crucial geometry of the ground appears. The prioritized processing of the ground geometry is consistent with our neuroimaging results, where subjects maintained central fixation, but OPA exhibited a more prominent representation of the floor (compared to the ceiling and the left and right walls; Fig. 8B).

## DISCUSSION

Visual navigation requires that the human brain represent the geometry of the environment, and especially the physical bounds of navigable space (Spelke and Lee, 2012). We refer to the stable geometric shape of the space bounded by walls, floor and the ceiling as the layout of the scene. Our results suggest that the scene-bounding elements are extracted from the visual input by the scene-responsive cortical area OPA. Specifically, using fMRI, we found that OPA encodes scene layout in a format that supports linear readout of the presence or absence of bounding elements with invariance to manipulations of surface texture. The representation in V1 was better captured by a model of low-level image features of the scene stimuli than the presence of the walls, whereas in OPA, the presence of the walls captured the representation better. PPA did not reliably distinguish the scene layouts in its fMRI response patterns, and overall, PPA showed better decoding performance for scene texture than layout, which was opposite to the results found in OPA. Texture is closely related to the category of the scene (e.g. indoor versus outdoor, urban versus natural). Our results hence support the view that OPA is involved in extracting the spatial structure of the local environment, whereas PPA has a role in scene recognition (Julian et al., 2018). We complemented the fMRI results with high-temporal-resolution MEG data. Scene-layout representations with similar invariance to surface texture as found in OPA with fMRI emerged rapidly after stimulus onset in the MEG responses (peaking at ~100 ms). The rapid emergence of an explicit encoding of scene layout in human OPA is consistent with recent modeling work suggesting that a layout representation can be efficiently computed by a feedforward mechanism (Bonner and Epstein, 2018).

Our stimulus set systematically varied the presence of five scene-bounding elements (three walls, floor, and ceiling) for a total of 2^5^ = 32 environmental geometries, each of which was rendered in three different surface textures. This systematic set enabled us to investigate the relative contribution of each of the five scene-bounding elements to the representations. The back wall covered a much larger, and more central, part of the visual field and thus mapped onto a larger patch of cortical surface in V1 (Duncan and Boynton, 2003) than other scene-bounding elements. The back wall was also prominently represented in downstream scene-responsive regions, which might reflect both its early visual representation and its navigational relevance in limiting forward movement. The removal of the back wall also adds depth to the scenes but the current stimuli were not optimised for fully disentangling the contribution of perceived depth from confounding low-level image features. Lescroart et al. (2015) have previously looked at the relationship between Fourier power and subjective distance in natural scenes. They conclude that these models provide similar predictions of brain activity in OPA and PPA when the responses to natural scenes are analysed using voxel-wise encoding models and suggest that some previous fMRI studies reporting representations of distance in scene-responsive cortical areas might be affected by this confound. More recently, Lescroart et al. (2019) used computer-generated stimuli similar to ours to model the relative contributions of low-level features and 3D structure on the voxel responses to scenes across the visual cortex. They report that individual voxels in scene-responsive areas represent combinations of orientations and distances of surfaces and, overall, that distance and openness are represented in OPA, PPA and RSC. Our results complement their results by showing a dissociation between the representations in OPA and PPA, with only OPA exhibiting a texture-invariant representation of scene layout, and by revealing the rapid emergence of this representation in the MEG responses.

The right and left wall, the ceiling and the floor all covered equal-sized regions in the visual field in our stimulus set, and hence, enabled comparisons of their relative contributions to the representations. The ceiling and floor differed from the left and right walls in terms of their prominence in the cortical representations. In V1, the ceiling was more weakly represented than the other scene elements, which is likely explained by the asymmetry between the upper and lower visual field representations in V1 (Henriksson et al., 2012; Liu et al., 2006). Overall, however, low-level image feature differences between the stimulus images better captured the representation in V1 than any scene-layout-based model. OPA exhibited a more prominent representation of the floor than of the left and right walls and the ceiling. One interpretation of this result is that the ground ahead of us has a special status in the OPA representation because it supports our movement in the environment. A special status for the ground is also supported by the eye-gaze data, collected before the neuroimaging experiments, which showed that subjects paid more attention to the floor than to the ceiling when viewing the novel scenes.

Consistent with our findings, previous studies have shown that OPA (also named transverse occipital sulcus, TOS) has lower- and peripheral-visual-field biases (Silson et al., 2015; Levy et al., 2004, 2001). An important question for future work is to what extent OPA represents the geometry of the ground ahead or more general features of the lower visual field. The OPA representation most likely reflects the natural regularities of the environment and the functional importance of different features. During normal navigation gaze is typically directed ahead, and hence, the floor is perceived in the lower peripheral visual field from where information is sampled to avoid obstacles and to adjust steps according to the terrain (Marigold and Patla, 2008; Turano et al., 2001). We have previously proposed that the occipital face area (OFA) might develop from a perifoveal retinotopic protomap into a map of face-feature detectors arranged on the cortical sheet in the topology of a face (the “faciotopy” hypothesis; Henriksson et al., 2015). The neighboring scene-responsive cortex has a peripheral visual-field bias (Levy et al., 2001, 2004) and might similarly develop from a retinotopic protomap into a representation of scene layout, with extended representations of the behaviourally most relevant features.

A limitation of the current study is the use 2D renderings of the scenes instead of 3D stimulus presentation. The literature on navigation has used virtual reality (VR) much more rigorously (for an example, see Hartley et al., 2003) than studies on visual perception, although scene perception studies would also benefit from 3D stimuli and interaction with the scenes. Challenges with more complex stimuli are stimulus control and how to reliably analyze and interpret the data. In the future, combining MEG with VR might allow novel experiments aimed at understanding the smooth interplay between visual perception and navigation during natural behavior. Furthermore, the scenes that we used as the stimuli likely varied in how plausible they are in the context of environments that we naturally encounter. We wanted to use a full stimulus set where the presence (absence) of each wall was equally possible. Future studies could reveal how cortical representations differ between scene layouts that are typically encountered and layouts that are improbable to encounter. Moreover, the use of a more restricted set of layouts with a wider set of textures could further help tease apart perceptual differences that in our stimuli might co-vary with the change in the layout, for example, related to perceived distance in a scene.

The functional roles of scene-responsive cortical visual areas OPA, PPA and retrosplenial cortex (RSC; Maguire, 2001) are often discussed in the context of navigation, but the interplay between visual perception and navigation is not yet fully understood. Navigation is thought to rely on a cognitive-map-like representation of the environment in the hippocampus that might be anchored to the visually perceived environment with the help of landmarks by the RSC (Epstein et al., 2017). Though PPA was originally linked to the perception of the spatial layout of the environment (Epstein and Kanwisher, 1998), cumulative evidence suggests that PPA is primarily involved in recognizing the scene category (e.g. kitchen versus woods) and scene identity (e.g. our kitchen versus someone else’s) using a combination of geometry and texture cues, whereas OPA is involved in the analysis of the local scene elements (Bonner and Epstein, 2017; Julian et al., 2018; Kamps et al., 2016; Lowe et al., 2017). Scenes that we recognize as distinct places might have identical layout and it is the textures and object ensembles that provide the crucial cues to where we are. Scene perception mostly happens in the peripheral visual field, which is limited by visual acuity but especially by visual crowding (Rosenholtz, 2016). PPA may extract the identity of the scene by encoding the summary statistics of object ensembles present in the scene (Cant and Xu, 2012). Overall, our results on the better texture than layout encoding in the PPA are in agreement with previous studies showing texture information in human PPA (Park and Park, 2017) and in the putative PPA homologue in macaques (Kornblith et al., 2013).

In conclusion, our study shows a striking distinction between PPA, which did not enable reliable decoding of scene layout, and OPA, whose activity patterns reflected the presence or absence of each of the five spatial constraints with invariance to surface texture. These results support the view that the OPA extracts information about a scene’s spatial structure, whereas PPA is involved in recognizing the scene context (Julian et al., 2018). Our analyses reveal the emergence, within 100 ms of stimulus onset, of a detailed encoding of scene layout in OPA, suggesting that this cortical region supports our rapid visual sense of the geometry of our environment.

## Acknowledgements

This work was supported by the Academy of Finland Postdoctoral Researcher Grant (278957) to LH, a British Academy Postdoctoral Fellowship (PS140117) to MM, and a European Research Council Starting Grant (ERC-2010-StG 261352) to NK. We thank ANI staff (especially Marita Kattelus, Mia Illman and Veli-Matti Saarinen) for assistance with the measurements. We acknowledge the computational resources provided by the Aalto Science-IT project.

## Author Contributions

Conceptualization, L.H., M.M., N.K.; Methodology, L.H., M.M., N.K.; Software, L.H., N.K.; Validation, L.H.; Formal Analysis, L.H.; Investigation, L.H.; Resources, L.H.; Data Curation, L.H.; Writing – Original Draft Preparation, L.H., M.M., N.K.; Writing – Review & Editing, L.H., M.M., N.K.; Visualization, L.H.; Supervision, L.H., N.K.; Project Administration, L.H.; Funding Acquisition, L.H.

## METHODS

### EXPERIMENTAL MODEL AND SUBJECT DETAILS

#### Subjects

Twenty-two healthy volunteers (9 females; mean age: 26, age range 19–49) with normal vision took part in this study. All subjects participated in a behavioural eye-tracking experiment, an MEG experiment and an fMRI experiment (divided into two sessions). Two additional subjects were recruited but excluded from the analyses because they did not participate in both fMRI and MEG experiments. Ethical approval for the research was obtained from Aalto University Ethics Committee. Subjects gave written informed consent before participating in the study.

### METHOD DETAILS

#### Spatial layout stimuli and experimental designs

We created all possible combinations of spatial layouts that can be made from five scene-bounding elements (left wall, back wall, right wall, floor, ceiling; 5^2^=32 different layouts). In practice, the layouts were created by starting from a 5-wall room and switching the spatial constraints ‘on’ and ‘off’ using the Blender 3D modelling software. Three different surface-textures and background images were applied. The textures were jittered for all stimuli to reduce the effect of low-level image similarities between the stimulus images. All 96 spatial layout stimuli are shown in Figure 1E.

In all experimental setups, the spatial layout stimuli extended approximately 20 deg × 20 deg of the visual field. The timing of the stimuli was controlled with Presentation^™^ (Neurobehavioral Systems, Albany, CA) software. In the behavioural lab, the stimuli were shown using a monitor. During the neuroimaging experiments, the stimuli were presented to semitransparent screens with 3-DLP projectors (Panasonic PT-D7700E; Panasonic PT-DZ110XEJ).

During the behavioural eye-gaze tracking experiment, the spatial layout stimuli were displayed for one second with a one second inter-stimulus-interval (ISI) when only a fixation cross was shown. The stimuli were shown once in a random presentation order that was different for each subject. The subjects were requested to return their gaze to the fixation cross in-between the stimulus presentations but no other instructions were given except that the subjects could freely explore the novel scenes.

The fMRI experiments applied a rapid event-related design, where the stimulus images were shown for 2 seconds followed by a 2-second ISI when only a red fixation cross was shown on a mid-grey background (stimulus trial). All 96 stimuli were shown once in a random order during one experimental run. The subjects were instructed to keep their eyes at the fixation cross throughout the experimental runs. To have enough baseline data, 32 4-second rest trials showing only the fixation cross were intermixed with the stimulus trials. Moreover, 10 task trials were randomly intermixed after the stimulus trials. During the task trials, an arrow was pointing to one of five directions and the subjects’ task was to respond with a button press whether there was a wall present in that direction in the previous layout. In sum, the total duration of a run was (96+32+10) * 4 seconds = 552 seconds, completed with a few extra rest-trials at the beginning and end of each run. Altogether 12 fMRI runs (12 trials for each stimulus) were collected for each subject. The fMRI data were collected during two separate experimental sessions.

During the MEG recordings, the stimulus images were displayed for 1 second followed by a 2-second ISI when only a red fixation cross was shown on a mid-grey background (stimulus trial). Similar to the fMRI experiments, 10 task-trials were randomly intermixed after the stimulus trials. All 96 stimuli were shown once in random order during one experimental run, and the subjects were instructed to keep their eyes at the fixation cross throughout the experimental runs. The total duration of one MEG run was (96 + 10) * 3 seconds = 318 seconds, completed with some baseline data at the beginning and end of each run. Altogether 8 MEG runs (8 trials for each stimulus) were collected for each subject.

#### Eye-gaze data collection and analysis

Eye-gaze data were collected separately in a behavioural laboratory and during the neuroimaging data collections. In all setups, we used SR Research EyeLink1000 systems (SR-Research Ltd., Ontario, Canada; sampling rate 500 Hz). A 9-point calibration was performed at the start of each experiment. During the fMRI and MEG experiments, the eye-gaze was tracked online to track the stability of the subjects’ gaze during central fixation. In the behavioural experiment, the subjects’ scan paths during free-viewing of the stimuli were followed. Fixations, saccades, and blinks were extracted from the continuous eye-tracking data using the software provided by the eye-tracker manufacturer.

#### (f)MRI data acquisition

The functional and anatomical MRI data were acquired using a 3T MAGNETOM Skyra whole-body scanner (Siemens Healthcare, Erlangen, Germany) equipped with a 30-channel head coil. During each main experimental run, the functional volumes were acquired using an EPI sequence with imaging parameters: repetition time 2 s, 29 slices with 2 mm slice thickness (no gap), field of view 192 mm × 192 mm, imaging matrix 96 × 96, echo time 30 ms, and flip angle 70°. Each subject attended two measurement sessions with six main experimental runs and two functional localizer runs. Two T1-weighted high-resolution structural images were acquired using an MPRAGE sequence, from which the white and gray matter borders were segmented and reconstructed using Freesurfer software package (Dale et al., 1999).

#### MEG data acquisition

MEG was recorded in a magnetically shielded room with a whole-scalp 306-channel MEG device (Elekta Oy, Helsinki, Finland). The device comprises 102 triple-sensor elements, with one magnetometer and two orthogonal planar gradiometers at each location. Only the gradiometer data (204 sensors) were used in the analysis here. The recording passband was 0.03–330 Hz, and the signals were sampled at 1000 Hz. The position of the subject’s head with respect to the MEG sensors was tracked throughout the experiment using four head position indicator coils. Horizontal and vertical electro-oculograms were recorded with the same recording passband and sampling rate as applied for the MEG.

### QUANTIFICATION AND STATISTICAL ANALYSIS

#### fMRI data analysis

Functional MRI data were pre-processed with SPM12 (Wellcome Department of Imaging Neuroscience) Matlab toolbox. The first four functional images from each run were excluded from the analysis to reach stable magnetization. The functional images were corrected for interleaved acquisition order and for head motion, and spatially smoothed using a 4 mm Gaussian smoothing kernel. The data from the second measurement session were co-registred and re-sampled to the same space with the first measurement session data. All analyses were performed in native space; no normalization was applied. The responses for the spatial layout stimuli were estimated using standard general linear model (GLM) analysis. The timings of the stimuli were entered as regressors-of-interest to the GLM, which were convolved with the canonical hemodynamic response model. Additional regressors included the responses and the six head-motion-parameters. During the parameter estimation, the data were high-pass filtered with 128-s cut-off.

The primary visual cortex (V1) was localized in each individual based on the cortical folds via a surface-based atlas alignment approach using Freesurfer (Hinds et al., 2008). Occipital place area (OPA), parahippocampal place area (PPA) and retrosplenial cortex (RSC) were localized based on independent functional localizer data. During the functional localizer run, the subjects were presented with blocks of images of scenes (different from the main experiment), faces, objects, and scrambled textures. Subjects performed a one-back task on the stimulus images. Two approximately 5-minute localizer runs were collected in both fMRI sessions for each subject. The regions-of-interest (ROIs) were manually drawn on each participant’s cortical surface based on the contrast scenes > faces using Freesurfer. Voxels representing the ROIs in the right and left hemispheres were concatenated for the linear discriminant analysis.

We used linear discriminant analysis (Kriegeskorte et al., 2007; Nili et al., 2014) to study the discriminability of the response patterns evoked by the different spatial layouts. The data were first divided into two independent sets based on runs. For each pair of spatial layout stimuli, Fisher linear discriminant analysis (Nili et al., 2014) was applied to find the weights for the voxels that discriminated between the response patterns and then the weights were applied to the independent data to calculate the linear-disciminant t-value (LD*t*), reflecting the discriminability between the response patterns evoked by two different spatial layouts. The LD*t* can be interpreted as a cross-validated, normalized version of the Mahalanobis distance (Nili et al., 2014). The analyses were done on individual data, and the linear-discriminant t-values were pooled across the 22 subjects. The significance was tested using a two-sided signed-rank test across the subjects. All pairwise comparisons of the spatial layouts were collected to matrices; multiple testing (496 pairwise comparisons of 32 spatial layouts) was accounted for by controlling the false-discovery rate. To test for generalization across surface textures, the Fisher linear discriminant was fit to the response patterns evoked by the spatial layouts in one texture and tested on the response patterns evoked by the same spatial layouts in another texture.

#### MEG data analysis

The continuous MEG data were preprocessed using spatiotemporal signal-space separation (Taulu and Simola, 2006) implemented in the MaxFilter software (Elekta Oy, Helsinki, Finland). This step included suppression of magnetic interference of external sources and compensation for head movement. Next, independent component analysis-based eye blink artifact correction was applied to the data (Oostenveld et al., 2010). Single-trial MEG responses to stimulus images were extracted from the continuous MEG recording, baseline-corrected from −200 ms to 0 ms and low-pass filtered at 45 Hz using tools provided by the MNE and Fieldtrip software packages (Gramfort et al., 2014; Oostenveld et al., 2010).

The cross-validated (leave-one-trial-out) Mahalanobis distance between the MEG responses was used as the distance estimator when constructing the representational dissimilarity matrices (RDMs; Walther et al., 2016). The RDMs were constructed separately for each time point using data from all 204 gradiometers.

#### Representational similarity analysis and fitting of models

For the fMRI data, the representational geometry (Kriegeskorte and Kievit, 2013; Kriegeskorte et al., 2008) of the regions-of-interest (V1, OPA and PPA) was characterized by the LD*t*-matrices. For the MEG data, cross-validated Mahalanobis distance was used as the distance estimator between the response-patterns of different stimuli. The representations were compared via the matrices using Kendall’s tau-a rank correlation (Nili et al., 2014). We used non-negative least-squares (Khaligh-Razavi and Kriegeskorte, 2014) (1) to fit multiple fMRI-LD*t*-matrices to an MEG distance matrix, and (2) to fit multiple competing models to brain representations (both fMRI and MEG). The fitting was done on individual distance matrices and the results were cross-validated using a leave-one-subject-out approach. In the cross-validation, the weights fitted separately for each subject were averaged across all subjects except the left-out subject. The left-out subject’s RDM was then compared to the reference-RDM with the average weights for the components. This approach was repeated by leaving out each subject in turn, and the results were averaged. The fitted models included the GIST (Oliva and Torralba, 2001) to capture low-level image features of the spatial layout stimuli and feature models based on the presence of different scene-bounding elements in the spatial layouts (see Fig. 6). The unique variance explained by each component model was evaluated by leaving out each model component in turn and evaluating the proportion of variance explained by the full model and the reduced model. The analysis was done on individual data and the results were averaged across subjects.

## SUPPLEMENTAL INFORMATION

**Supplementary Figure S1.**
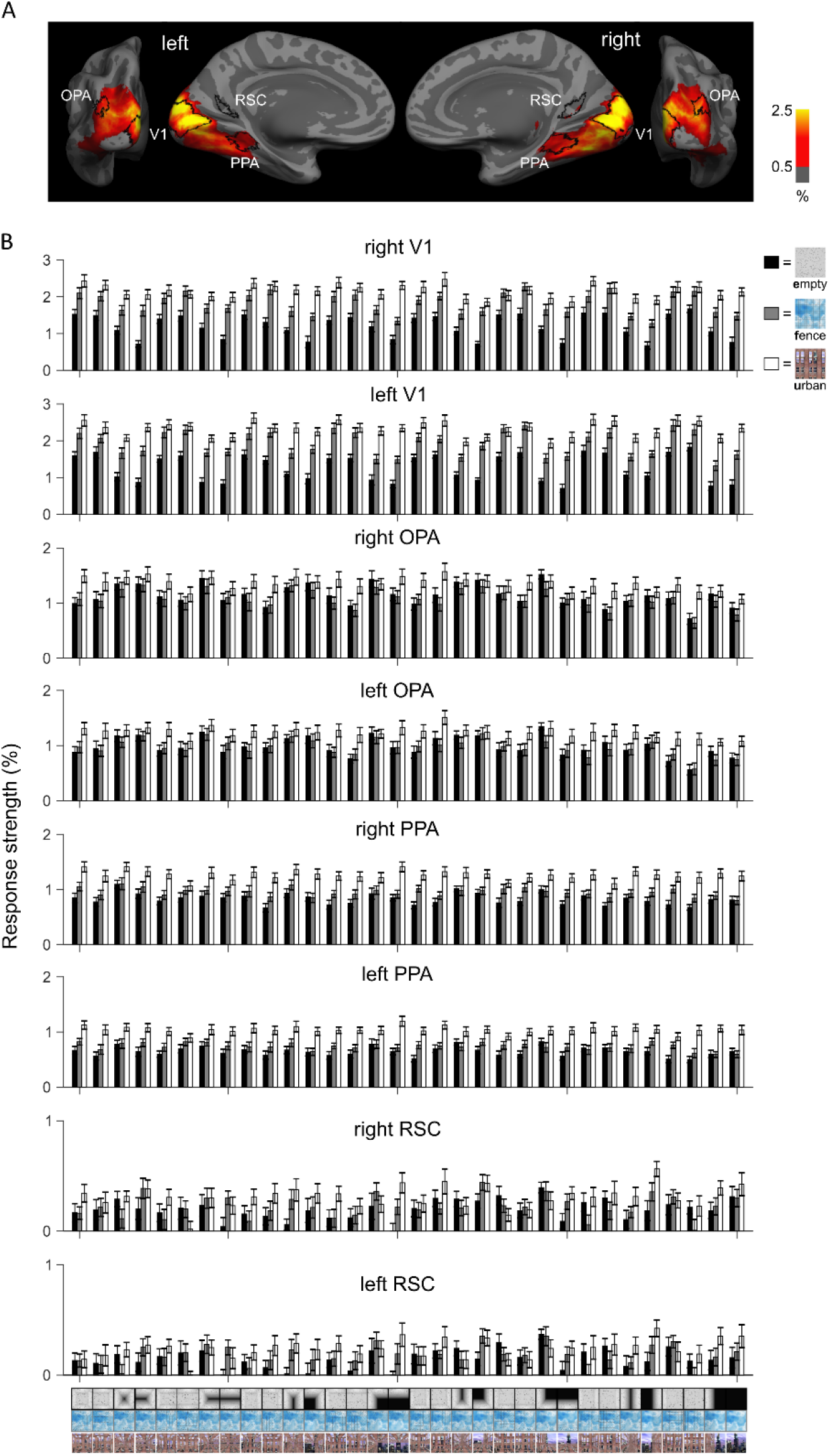
Average fMRI responses to the 96 scene stimuli (related to Figure 2). A) Average response to the scene stimuli is shown on the group-average cortical surface. The average locations of the regions-of-interest (V1, OPA, PPA, RSC) are also shown. The response strength (%) for each stimulus was calculated from the beta weight for the stimulus divided by the average beta weight for the constant terms (across runs) multiplied by 100. The responses were averaged across all stimuli and subjects. The map is thresholded at 0.5%. B) Average response to each scene stimulus is shown separately for each region-of-interest. Scenes with different textures are shown in different colours (black, grey, white). The error bars indicate standard errors of the mean (SEMs) across the 22 subjects.

**Supplementary Figure S2.**
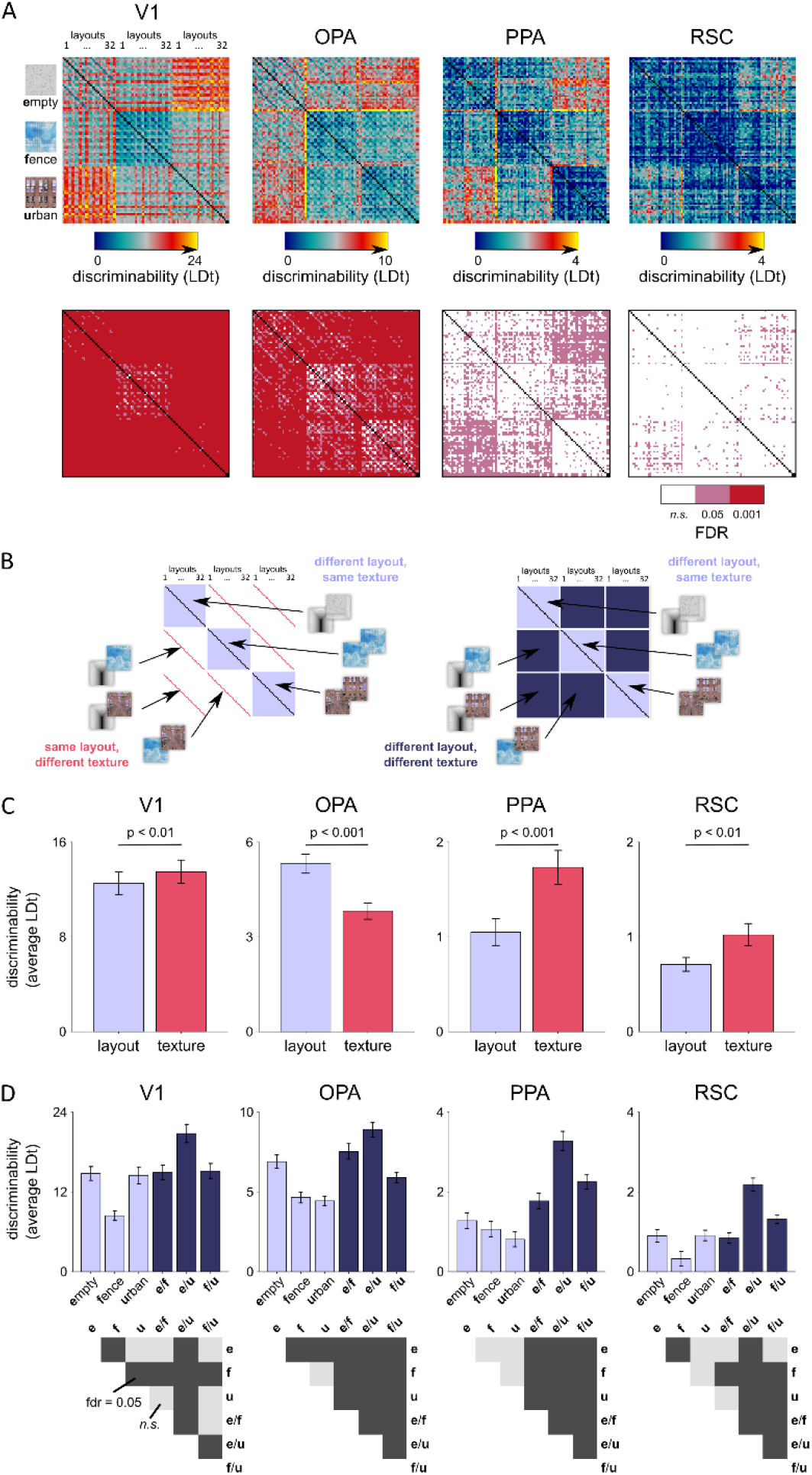
Discriminability of the fMRI response-patterns for each pair of scene stimuli (related to Figure 2). A) The discriminability of the scene stimuli from the fMRI response-patterns was evaluated for each pair of the 96 scenes. The Fisher linear discriminant was fitted to the response-patterns obtained from the training data (odd fMRI runs) and the performance was evaluated on independent testing data (even fMRI runs). The results are shown as linear discriminant t-values (LDt). The analyses were done on individual data and the results were pooled across the subjects. B) The average discriminability was calculated for (1) scene pairs that had a different layout but same texture, (2) scene pairs that had same layout but different texture, and (3) scene pairs that had different layout and different texture. C) In V1, PPA and RSC, the average discriminability was higher between scenes that had a different texture than a different layout. In OPA, the average discriminability was higher between scenes that had a different layout than a different texture. The p-values were obtained using a two-sided signed-rank test across 22 subjects. The error bars indicate standard errors of the mean (SEMs) across the 22 subjects. D) The average discriminabilities are shown separately for each texture (scenes differ in layout; light purple) and texture pairs (both the texture and layout are different between the scenes; dark purple). Only in PPA, the average discriminability was consistently higher when both the scene layout and the scene texture was different between the pair of scenes (dark purple bars) compared to when the scenes only differed in their layouts (light purple bars). The error bars indicate standard errors of the mean (SEMs) across the 22 subjects. The bottom row shows the expected false-discovery rates (FDR; n.s., not significant); the p-values were obtained using a two-sided signed-rank test across 22 subjects.

**Supplementary Figure S3.**
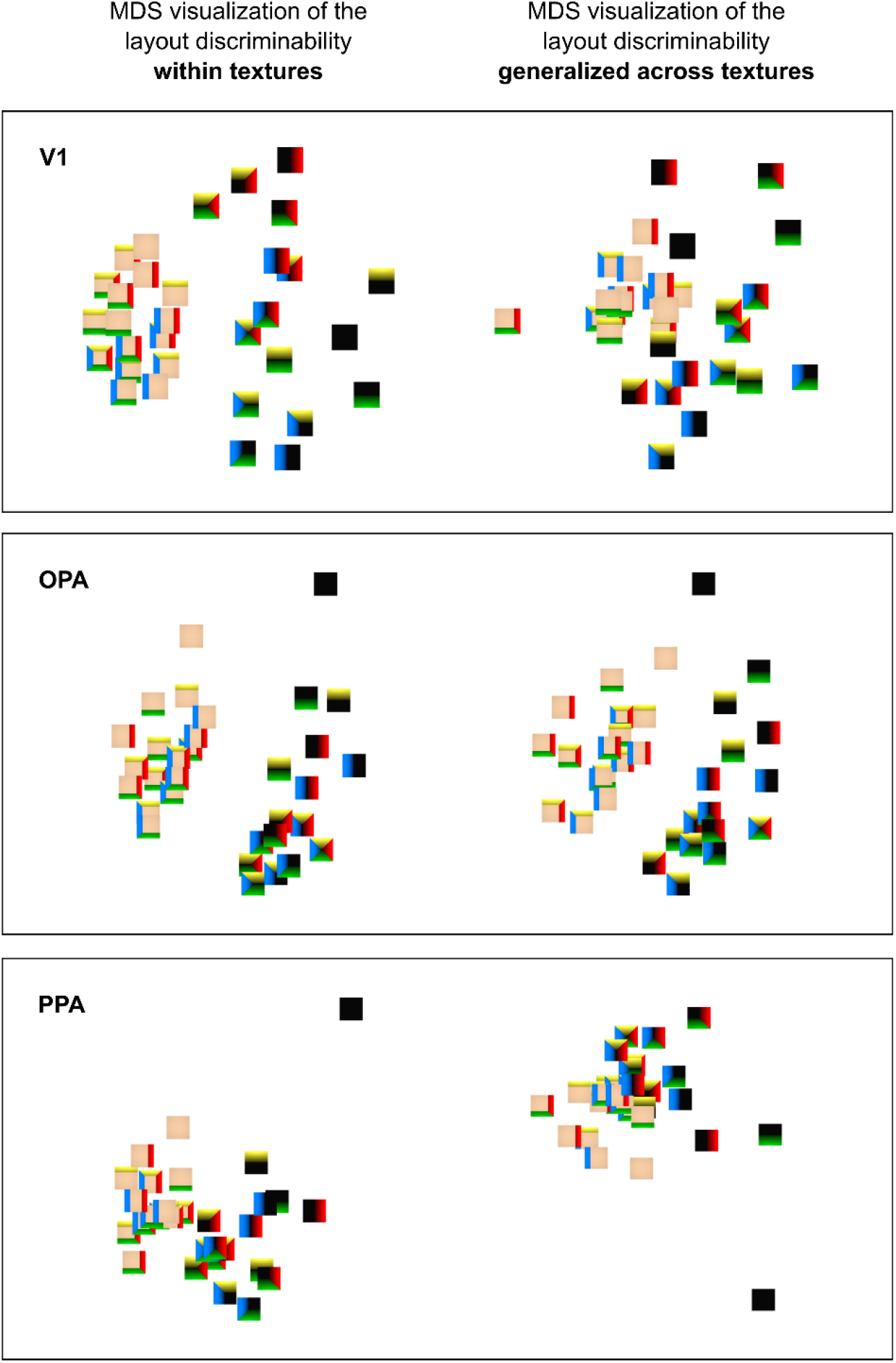
Multidimensional scaling visualization of the representational geometry in V1, OPA and PPA (related to Figure 4). The multidimensional-scaling (MDS; metric stress) visualizations of the distinctiveness of the response-patterns using simplified layouts (walls coded with different colours as in Fig. 1). On the left side the discriminability of the layouts was evaluated within textures (results averaged across the three textures; RDM shown in Fig. 4A). On the right side the generalization of the discriminability of the layouts across textures was evaluated (results averaged across the different combinations; RDM shown in Fig. 4B). The representation of scene layout shows a better generalization across textures in OPA than in V1 or PPA.

**Supplementary Figure S4.**
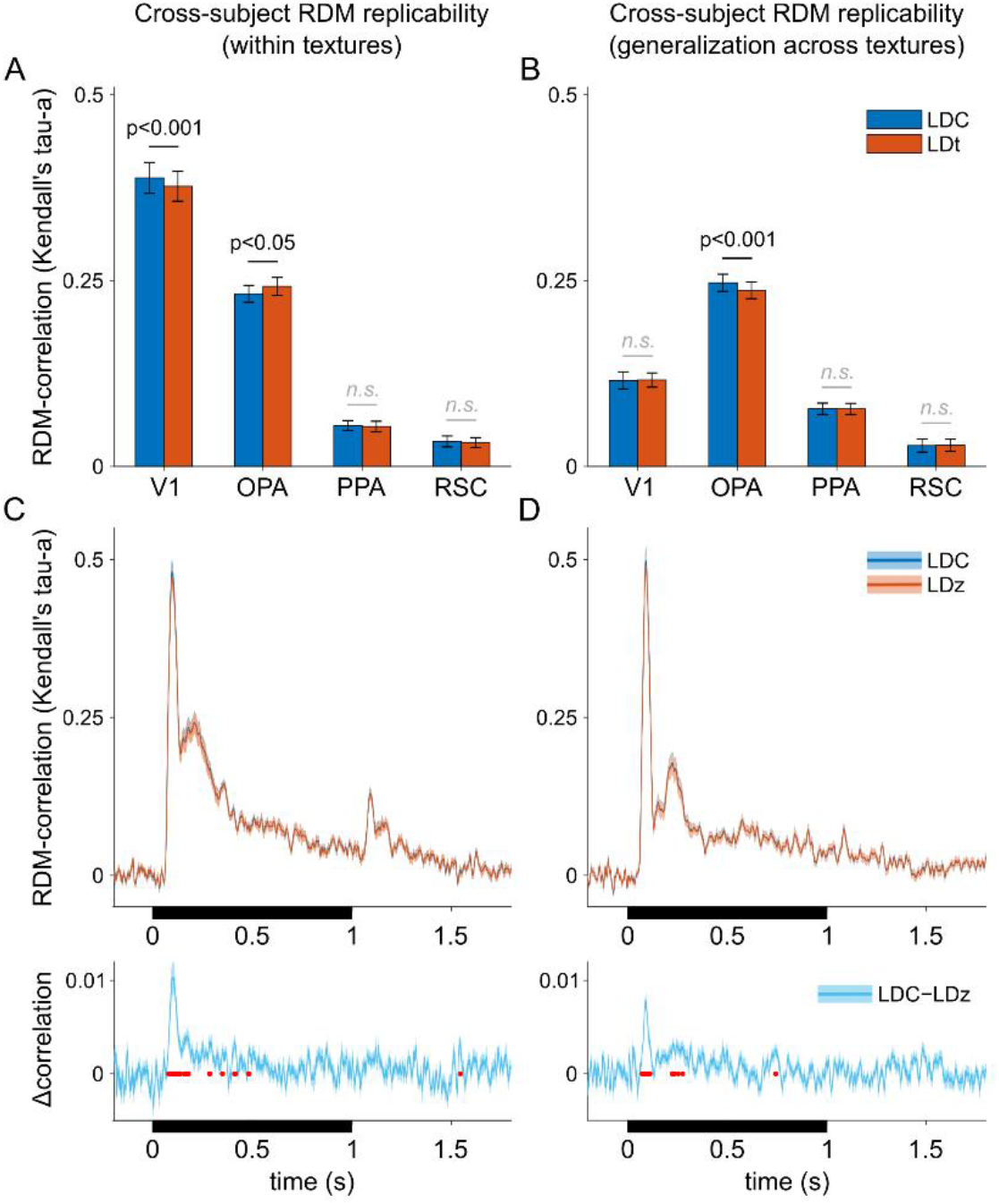
Effect of cross-validated distance estimator normalization on the cross-subject RDM replicability (related to Figure 5). A–B) The reliability of the RDMs was evaluated by assessing the RDM replicability across subjects (Kendall’s tau-a rank correlation). For fMRI data, the RDMs for each subject were constructed either using a cross-validated estimate of the Mahalanobis distance (linear discriminant contrast, LDC) or using the LDC normalized by an estimate of its standard error (linear-discriminant t value, LDt). The standard error was estimated from the residuals of the independent testing data after projection on the discriminant dimension (Nili et al., 2014). The error bars indicate standard errors of the mean (SEMs) across the 22 subjects. The p-values (n.s., not significant) were obtained using a two-sided signed-rank test across 22 subjects. C) For MEG data, the RDMs for each subject were constructed either using a cross-validated estimate of the Mahalanobis distance (linear discriminant contrast, LDC) or using the LDC normalized by the baseline period (linear-discriminant z value, LDz). Shaded regions indicate SEM across subjects, red line indicates false-discovery rate of 0.05 (two-tailed signed-rank test across the 22 subjects; FDR across time points).

**Supplementary Figure S5.**
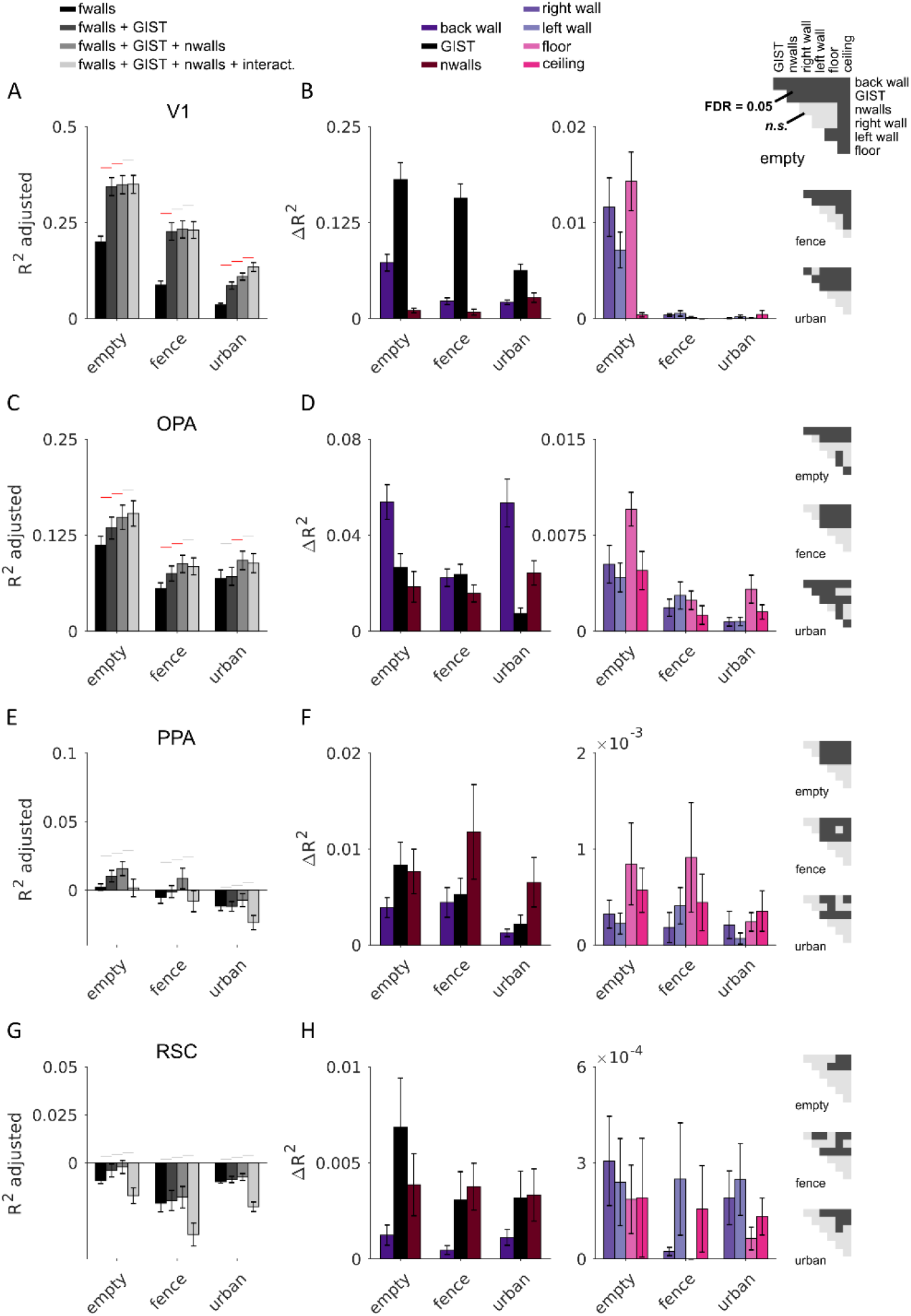
The unique variance in fMRI-RDMs explained by each model component, shown separately for each texture (related to Figure 8). The explained variance (R^2^ adjusted for the number of component models) of the fitted joint models containing different component models (1: component for each wall; 2: component for each wall and the GIST model; 3: component for each wall, GIST model and number of walls; 4: component for each wall, GIST, number of walls and interactions between the walls). The results are shown separately for each scene texture (empty, fence, urban) for V1 in (A), OPA in (C), PPA in (E), and RSC in (G). The red lines indicate a significant improvement compared to a model with fewer components (one-tailed signed-rank test across the 22 subjects, multiple testing accounted by controlling the false discovery rate at .05; the grey lines indicate a non-significant difference). The gain in the variance explained by each of the model components was calculated by comparing the explained variance between the full model (3: component for each wall, GIST model and number of walls) with a model where this one component was left out from the joint fit. Each component was left out in turn, and the results are shown separately for each texture and for V1 in (B), OPA in (D), PPA in (F), and RSC in (H). In different regions-of-interest, the different model components made a different contribution to the representation; see the half-matrices for results from the significance testing (two-tailed signed-rank test across the 22 subjects, multiple testing within an analysis accounted for by controlling the false discovery rate at .05).

**Supplementary Figure S6.**
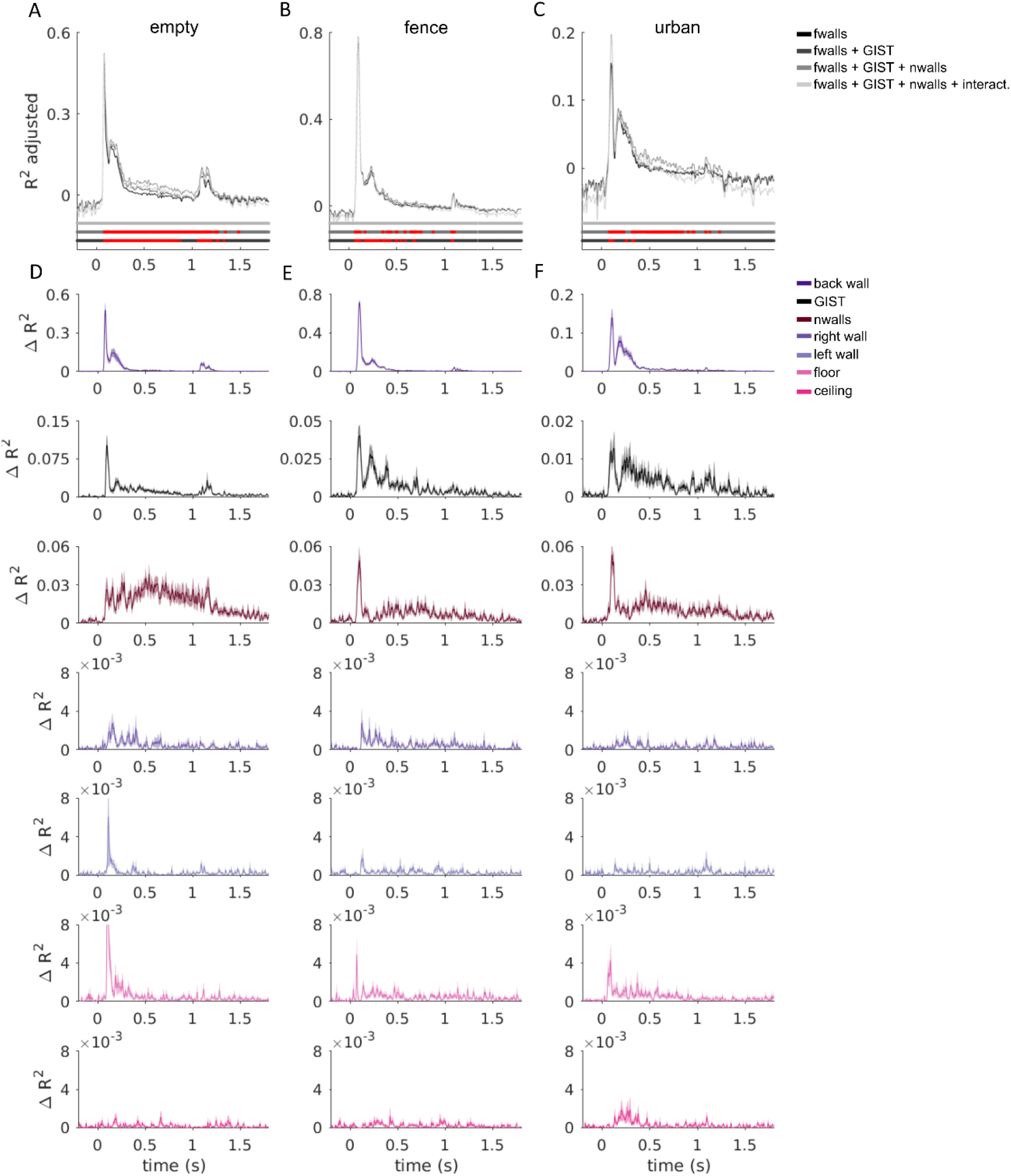
The unique variance in MEG-RDMs explained by each model component, shown separately for each texture (related to Figure 8). The explained variance (R^2^ adjusted for the number of component models) of the fitted joint models containing different component models (1: component for each wall; 2: component for each wall and the GIST model; 3: component for each wall, GIST model and number of walls; 4: component for each wall, GIST, number of walls and interactions between the walls). The results are shown separately for each scene texture in A–C. The red lines indicate a significant improvement compared to a model with fewer components (one-tailed signed-rank test across the 22 subjects, multiple testing across time-points accounted by controlling the false discovery rate at .05; the grey lines indicate a non-significant difference). The gain in the variance explained by each of the model components was calculated by comparing the explained variance between the full model (3: component for each wall, GIST model and number of walls) with a model where this one component was left out from the joint fit. Each component was left out in turn and the results are shown in different rows using the same colours as in Fig. S5 for fMRI data.

**Supplementary Figure S7.**
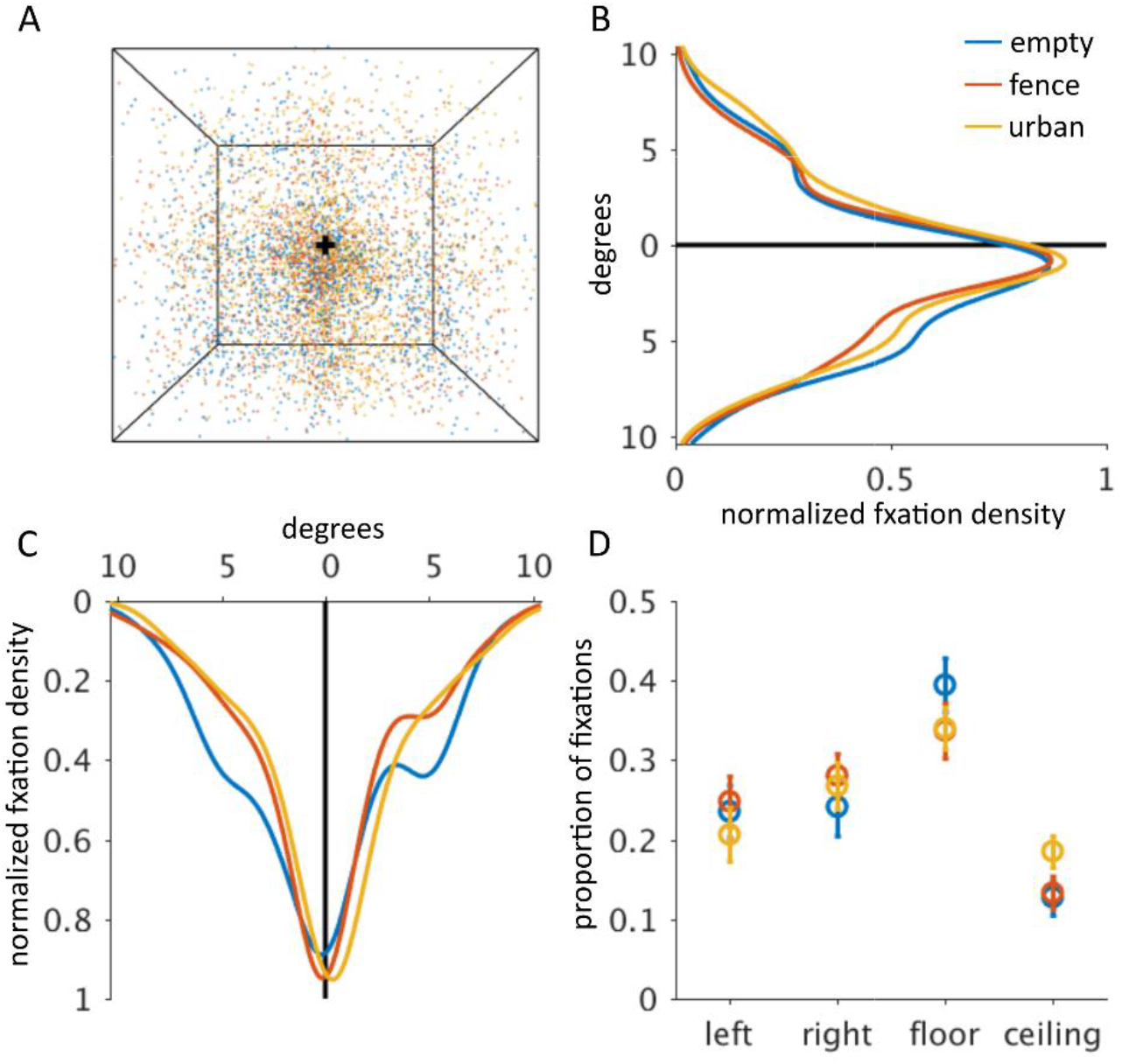
Gaze allocation during free-viewing of the spatial layout stimuli highlights the lower parts of the scenes (related to Figure 8). A) Before the neuroimaging experiments, subjects’ gaze during free-viewing of the spatial layout stimuli were collected. All individual fixations for all of the stimuli (Fig. 1E) are shown; the different colours represent the three different surface textures (Fig. 1D). The start position was always at the centre of the stimuli (shown with black cross). B) Gaze was more often directed to the lower than to the upper part of the scene. C) The left and right parts of the scenes were looked at equally. D) The proportion of fixations was greater to the floor than to the ceiling (FDR = 0.0035; 0.0077; 0.003 for each texture separately; two-tailed signed-rank test across 22 subjects; FDR across the combinations of wall pairs).

